# AlphaFold2-RAVE: From sequence to Boltzmann ensemble

**DOI:** 10.1101/2022.05.25.493365

**Authors:** Bodhi P. Vani, Akashnathan Aranganathan, Dedi Wang, Pratyush Tiwary

## Abstract

While AlphaFold2 is rapidly being adopted as a new standard in protein structure predictions, it is limited to single structure prediction. This can be insufficient for the inherently dynamic world of biomolecules. Even with recent modifications towards conformational diversity, AlphaFold2 is devoid of providing thermodynamically ranked conformations. AlphaFold2-RAVE is an efficient protocol using the structural outputs from AlphaFold2 as initializations for AI augmented molecular dynamics. These simulations result in Boltzmann ranked ensembles, which we demonstrate on different proteins.

---

While protein structure prediction has traditionally relied on different experimental techniques, 2021 saw a change in the status quo with AlphaFold2^*1*^. It has surpassed the accuracy of other models^*2,3*^ and offers a seemingly robust tool for structure prediction directly from amino acid sequences, relying largely on the idea that conservation of residues across evolutionary protein sequences is correlated with three-dimensional Euclidean distances. Building on this, AlphaFold2 generates multiple sequence alignments (MSA) of evolutionarily related sequences, hence identifying residues that co-evolve to facilitate structure prediction.

While AlphaFold2 indeed represents phenomenal leaps forward for structural biology, it has some key limitations^*4–6*^ with no clear solution so far. The first is that in its original incarnation, AlphaFold2 is a single structure prediction method. Biology is often not about a single structure, but about the ensemble of inter-converting structures^*7*^. This problem was first addressed by simply reducing the size of the input MSA in AlphaFold2, and effectively increasing the conformational diversity explored^*5,8*^. However, this does not provide any notion of relative probabilities of these conformations, even as it introduces several thermodynamically unstable or improbable structures. A second limitation is that AlphaFold2 fails in predicting changes in protein structure due to missense mutations^*4*^. Being able to assign Boltzmann weights would instantly rule out unphysical models generated by MSA length reduction, simulataneously ranking them by thermodynamic propensity, thereby addressing both limitations.

In this communication, we solve these limitations using Artificial Intelligence (AI) augmented all-atom resolution Molecular Dynamics (MD) methods^*9*^. In principle, long unbiased MD simulations, with no AI or AlphaFold2 assistance, should directly characterize the conformational diversity and thermodynamics of all proteins. However, MD simulations are limited in timescales, making sampling diverse protein conformations with statistical fidelity impossible^*10*^. We propose a protocol wherein starting from a given sequence, we obtain an ensemble of Boltzmann-weighted structures, i.e. structures with their thermodynamic stabilities. We combine AlphaFold2 in post-processing with the AI-augmented MD method “Reweighted Autoencoded Variational Bayes for Enhanced Sampling (RAVE)”^*9*^, calling the final protocol AlphaFold2-RAVE. RAVE is one of many enhanced MD methods that surmount the timescale limitation; in Methods we provide an overview, the advantages it provides over other enhanced MD methods, and other technical details.

We present illustrative results using AlphaFold2-RAVE on proteins with unique challenges: the 1HZB CSP^*11*^ for side-chain orientation predictions, ubiquitin binding protein UBA2 for disorder effects of missense mutations^*4,12–14*^, and SSTR2 GPCR for conformational diversity predictions^*15*^. For each, we show that AlphaFold2 fails, even with the reduced MSA trick from Ref. (*5*). AlphaFold2-RAVE does a perfect job in reproducing benchmarks for CSP known from experiments and specialized simulations, while providing results that correlate with both experimental results and biological roles for UBA2 and GPCR.

Our central idea is to first use reduced-MSA AlphaFold2 to generate many possible conformations as the initialization for RAVE, which uses an autoencoder-inspired framework to learn relevant slow degrees of freedom, also called reaction coordinates (RC), by iterating between rounds of MD and autoencoder based analysis, wherein every MD iteration is biased to enhance fluctuations along the new RC. The RC itself is expressed as a “State Predictive Information Bottleneck (SPIB)”^*16*^, i.e. the least information needed about a protein’s current attributes to predict its future state after a specified time. This allows one to account for the inherently dynamic personalities of proteins^*17*^, as well as obtain a Boltzmann-weighted ensemble of conformations.

Fig. 1 shows results for the 66-residue CSP (PDB:1HZB). CSP has known rotameric metastability in its eighth residue Trp8 (Fig. 1b), exhibiting 6 states in its *χ*_1_ and *χ*_2_ torsions through fluorescence spectroscopy^*11*^. To demonstrate the power of AlphaFold2-RAVE, we assume no prior knowledge of any such special residues. Fig. 1c show probability distributions of conformations obtained from AlphaFold2 with reduced MSA length and with AlphaFold2-RAVE respectively, projected in the space of SPIB coordinates. The enhancement in quality of sampling is unequivocally evident. Further analyses are provided in SI establishing that the enhanced sampling here conforms to the Boltzmann distribution, and that our protocol is also more general than focused sampling.

**FIG. 1:**
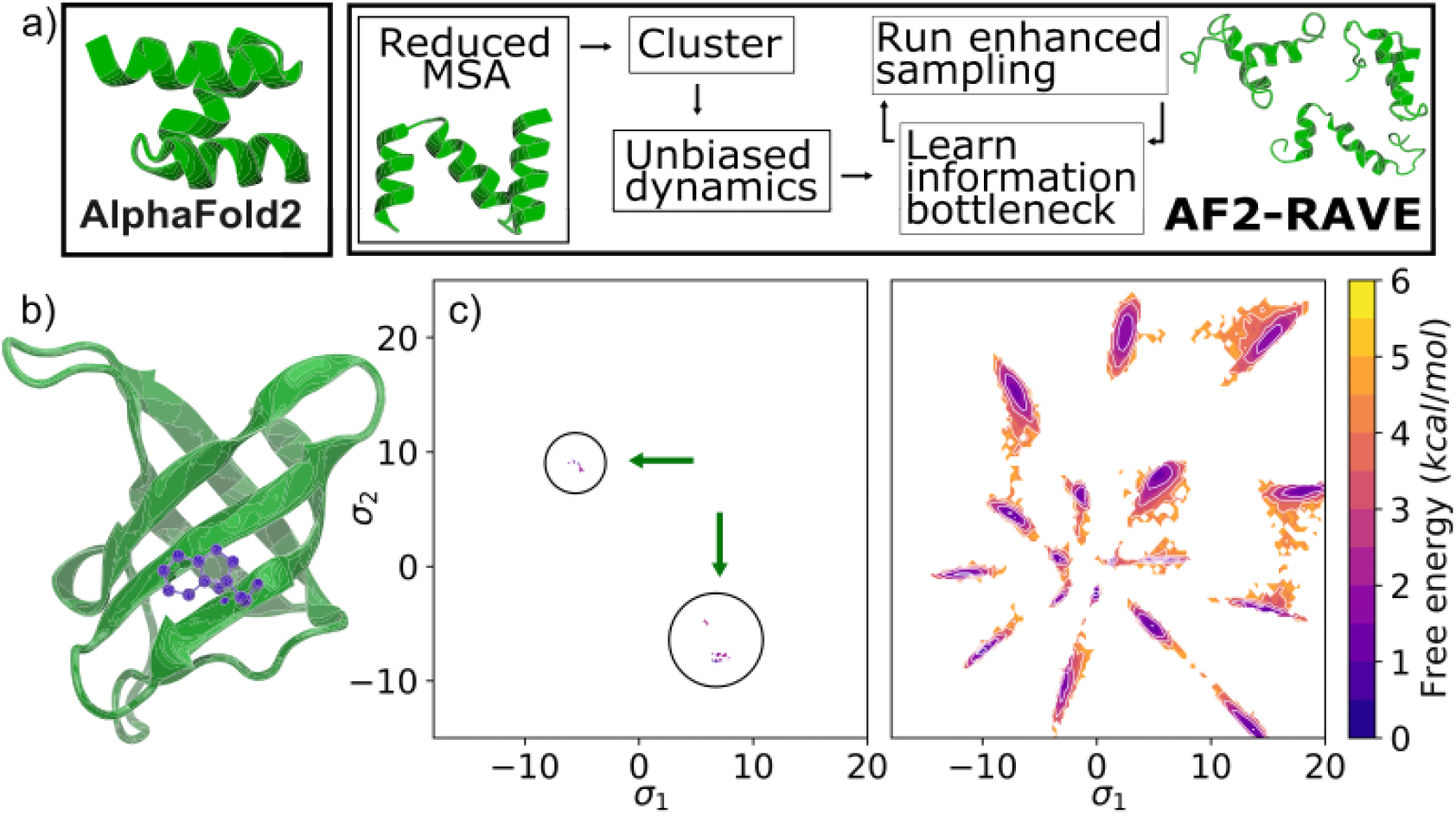
a) AlphaFold2-RAVE schematic with representative UBA2 L355A structures from AlphaFold2, reduced-MSA AlphaFold2, and partially disordered structure predicted by AlphaFold2-RAVE. b) CSP 1HZB structure with Trp8 in purple, c) Results using AlphaFold2-RAVE on CSP. Potentials of Mean Force (PMFs) projected along the two-dimensional SPIB *σ*_1_ and *σ*_2_ learnt from our scheme. Projections using different methods shown. Left: Distribution from reduced MSA showing some diversity compared with AlphaFold2, however without thermodynamic reliability. Right: Results from AlphaFold2-RAVE, showing both conformational diversity and thermodynamics.

Fig. 2c shows PMFs from AlphaFold2-RAVE for both WT and L355A UBA2. AlphaFold2 was recently shown to be unable to capture the changes in native state stability of a partially helical disordered structure caused by the missense mutation L355A^*4*^. For the WT we find that the folded-unfolded state energy difference is −0.3 kcal/mol, while for the mutant the same difference is 1.2 kcal/mol. This shows that the mutation does indeed increase its disordered nature. In Fig. 1 we show representative structures obtained from all three stages of our protocol, demonstrating our significantly enhanced sampling of quasi-helical disordered structures.

**FIG. 2:**
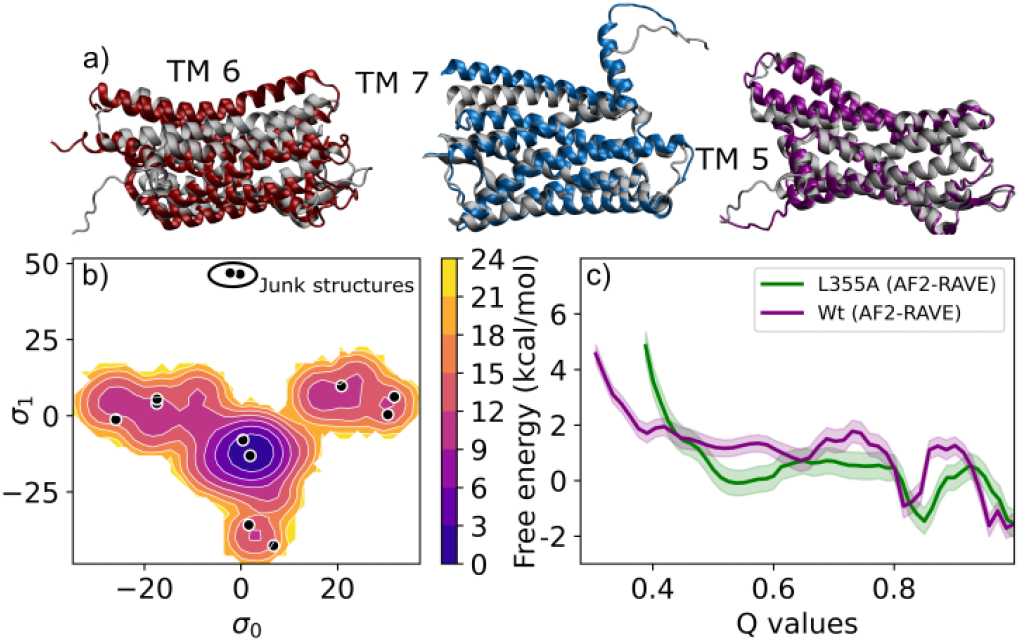
a) SSTR2 structures highlighting helix movement in TM6, TM7, TM5 via AlphaFold2-RAVE enhanced sampling, agreeing with experiments^*18*^ and biological function. Low energy structures are in colour, overlaid on native structure (grey). b) SSTR2 PMF using AlphaFold2-RAVE, showing multiple potential states. c) UBA2 PMFs along total Q-value for WT(purple), L355A(green). The wild type shows a visibly higher barrier and L355A has a wider, stabler disordered region.

The final system we study is the medically relevant^*19*^ G-protein-coupled receptor (GPCR), for which AlphaFold2 is unable to capture conformational diversity^*14*^. Specifically, we explore conformational shifts in the somatostatin receptor SSTR2. Once again, we find that AlphaFold2-RAVE detects several local changes in sidechain positions, and also larger scale helical motions. This corresponds well with known evidence of GPCR structural diversity^*18*^. In Fig. 2a, we show examples of the large scale helical shifts we are able to observe. Fig. 2b shows the PMF projected on the SPIB.

To conclude, we propose the AlphaFold2-RAVE method, combining strengths of AlphaFold2 with all-atom resolution enhanced sampling powers of RAVE^*9*^. This provides us with an ensemble of diverse conformations ranked with their correct Boltzmann weights. While any sampling protocol could in principle be implemented here, we prefer RAVE (specifically the SPIB variant) due to minimal hand-tuning and prior knowledge required regarding metastable states and latent spaces to drive sampling. Similarly, instead of AlphaFold2 we could use other experimental^*20*^ or computational^*2,3*^ approaches to generate initial dictionaries of possible conformations. We choose AlphaFold2 for its simplicity, accuracy and relative ease. Here, we apply AlphaFold2-RAVE to three systems of biological relevance, demonstrating its usefulness in obtaining conformational diversity.

Individual components of the code are available at github.com/sokrypton/ ColabFold and github.com/tiwarylab/ State-Predictive-Information-Bottleneck.

## Acknowledgments

Research in this publication was supported by the National Institute of General Medical Sciences of the National Institutes of Health under Award Number R35GM142719. The content is solely the responsibility of the authors and does not represent the official views of the National Institutes of Health. We thank Deepthought2, MARCC, and XSEDE^*21*^ (projects CHE180007P and CHE180027P) for computational resources. We thank Dr. Eric Beyerle, Zachary Smith, and Luke Evans for critical reading of the manuscript, and Dr. Ed Miller and Dr. Dilek Coskun for helpful discussions regarding GPCRs.

## METHODS

### A. AlphaFold2

To generate a diverse structural ensemble, we made a key change to the publicly available Colabfold notebook^*22*^. The input featurization for the AlphaFold2 algorithm is in the form of two MSA clusters. The first is used as the input to the main AlphaFold2 network, and the second is used to initialize a pair representation matrix, which is the input for a second channel in the AlphaFold2 network. We decrease the size of both MSA clusters to (16, 32) and (8, 16), to obtain local conformational changes in CSP and SSTR2. For UBA2, to get more global changes, we further decrease MSA length to (4, 8) and (2, 4). As also reported in Ref. (*5*), we find that restricting the information in this featurization leads to revealing structures that are significantly different from the crystal structure, while providing hints to metastability. The other computational choices and parameters are as follows. We use the model dropout feature, set number of recycles to 1, and use 128 random seeds– since AlphaFold2 generates 5 structures per random seed, we obtain 640 structures for each set of MSA hyperparameters.

### B. Metadynamics

Algorithms developed to enhance sampling can be characterized in two ways: driving sampling by adding “biases” in the form of energetic potentials, or by splitting and stratification methods that focus on simulating multiple trajectory fragments and resampling them with biased probabilities^*23*^. In practice, the former is found to be preferable for enthalpic barriers, while the latter is more effective for entropic barriers. Our scheme is in theory agnostic to the sampling method used, as long as it is possible to recover unbiased statistics to obtain potentials of mean forces using the method. Here, we choose to use an easily implementable and relatively fast method, metadynamics^*24*^. Metadynamics functions by depositing Gaussian biases intermittently along the simulation so that regions of the collective variable space that have been traversed become less probable the more they are sampled, and the system is forced to sample rarer regions. This results in a time dependant bias that is recorded and can be used to generate a PMF along the collective variables used for bias. Additionally, it has been shown that an on-the-fly bias can be computed independently^*25*^ to calculate a weighted histogram along any arbitrary collective variable. However, while essential, in practice the choice of collective variable is difficult. Biasing irrelevant collective variables or missing crucial slow collective variables^*26*^ could result in observing no significant motion away from initial metastable state, or to sampling unphysical regions of configurational space.

### C. State predictive information bottleneck

Here, we use SPIB iteratively with biased or enhanced dynamics to learn the reaction coordinate^*16*^. SPIB is essentially a past-future information bottleneck protocol which takes time resolved trajectory data in a high dimensional order parameter space. It then predicts a latent space of desired dimensionality and identifies states in this space. Here we used a 2-dimensional information bottleneck which we denote {*σ*_1_, *σ*_2_}. The protocol requires an initial identification of state labels which is then refined iteratively. The model learnt through SPIB consists of an encoder, which transforms the high dimensional input into a latent space, and a decoder, which uses this latent space to predict state labels after a preset time-lag. Ideally, the latent space identified as the information bottleneck would correspond with our traditional understanding of reaction coordinates. This was demonstrated to be true for a model system in Ref. (*16*), where the latent space corresponds very well with the committor as defined in transition path theory. The information bottleneck is found by optimizing a loss function that maximizes the latent space’s ability to predict the state of the system after a pre-set time-lag, while reducing the input dimensionality to the given bottleneck dimensionality. SPIB is capable of using both a linear or a non-linear encoder. In this work, we use a linear encoder with a non-linear decoder. For CSP, the encoder was a 2-d linear combination of 125 order parameters arising from the sidechain dihedral angles *χ*-s of all 66 residues. We have found that while there are regions of state space that have zero sampling that we may aim to push the simulation into, a linear encoder is far superior as a non-linear encoder will often overfit to sampled regions while producing unphysical results in unsampled regions.

### D. Other simulation details

The protein is represented by the AMBER03 force field^*27*^. The simulations are performed at 300 K with the leap-frog integrator in GROMACS 5.1.4^*28*^; LINCS was used to constrain the lengths of bonds to hydrogen atoms^*29*^; the step size was 2 fs. We used PLUMED 2.4^*30*^ to extract collective variables.

### E. Collective variable choices

An important feature of our protocol, both in its versatility and adaptability, is the choice of input order parameters or collective variables in the clustering and SPIB learning stages. On one hand, SPIB’s ability to handle a large number of inputs lends itself to highly exploratory work, the careful choice of CVs also lends itself to our ability to study extremely different scientific questions. In our work, we present three unique classes of protein structural changes, and our choice of CVs is key.

For CSP, we opt for the most generalized choice of collective variables to study rotameric states: all sidechain dihedral angles. For the 66-residue peptide, this results in 125 dihedral angles.

Similarly, for UBA2 to retain maximum information while exploring partially disordered states, we opt for the per residue Q value^*31*^ and inter-helical angles as our input collective variables, defined for a given structure as follows:

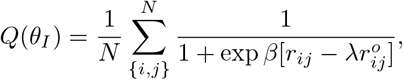

where *θ_I_* is the *I* th residue, *r_ij_* is the distance between atoms *i* and *j* for the specified structure, and 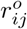 is the distance in the reference (native-like) structure. The atom pairs *i, j* are chosen so that *i* belongs to the residue *I*, and 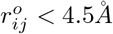, i.e. all pairs of atoms that are geometrically in “contact” in the reference structure. We choose the single structure Alphafold2 outputs as our reference both wild-type and mutant. The parameters *β* and *λ* were taken from Ref. (*31*). For a notion of inter-helical angles, we use the *α*-carbons of residues {R9, F15, I21} and {A23, K29, A35}.

Our approach for SSTR2 is more targeted, as we specifically aim to sample conformations that change ligand binding affinities, so as to facilitate drug discovery. SSTR2 has a recently resolved experimental structure, and like most other GPCRs, the residues involved in ligand binding are known^*32*^. We begin by narrowing our CVs to this set of residues, tabulated in Table 1, then selecting all residue pairs that are (1) within 4.5Å of each other and (2) do not belong to the same transmembrane helix or connecting loop. We use the C*α* distances between these residues as our initial OPs. One key difference in protocol is that to select initial structures for unbiased sampling, we use regular space clustering^*33*^, which results in a more diverse and less redundant initial guess, as AlphaFold2 tends to overamplify the probability of native-like or inactive structures.

**TABLE I:** Residues used to compute pairwise distances as inputs for RAVE.

|  |  |  |  |
| --- | --- | --- | --- |
| PHE92 | GLN102 | ASP122 | GLN126 |
| PHE127 | ARG184 | SER192 | THR194 |
| ILE195 | ASN196 | TRP197 | GLY199 |
| TYR205 | PHE208 | ILE209 | THR212 |
| PHE272 | PHE275 | ASN276 | PRO286 |
| LYS291 | PHE294 | TYR302 |  |

