## Supplementary Materials for "AlphaFold2-RAVE: From sequence to Boltzmann ensemble"

### Supplemental Information for
### AlphaFold2-RAVE: From sequence to Boltzmann ensemble

Bodhi P. Vani*

*Institute for Physical Science and Technology,*

*University of Maryland, College Park, Maryland 20742, USA*

Akashnathan Aranganathan*

*Biophysics Program and Institute for Physical Science and Technology,*

*University of Maryland, College Park 20742, USA*

Dedi Wang

*Biophysics Program and Institute for Physical Science and Technology,*

*University of Maryland, College Park 20742, USA*

Pratyush Tiwary†

*Department of Chemistry and Biochemistry and Institute for Physical Science and Technology,*

*University of Maryland, College Park 20742, USA*

(Dated: October 14, 2022)

* These two authors contributed equally.

**Appendix A: Benchmarking the protocol using cold shock protein**

To benchmark our protocol, we also run a control simulation on the cold shock protein (CSP; PDB ID: 1HZB). In this work, we present sampling on the State Predictive Information Bottleneck (SPIB), a linear combination of all sidechain dihedrals to show the existence of several rotameric states. Here, we ensure that our method both samples the correct thermodynamic profile for the dihedrals of the specific residue of interest, while also increasing ergodicity in sampling in different degrees of freedom.

CSP has known rotameric metastability in its eighth residue (Trp8), exhibiting 6 metastable states in its $\chi_1$ and $\chi_2$ (the first two dihedral angles following the backbone) torsions as per fluorescence spectroscopy [1]. We show representative structures for these 6 states with the relevant residue highlighted in purple in Fig. S1.

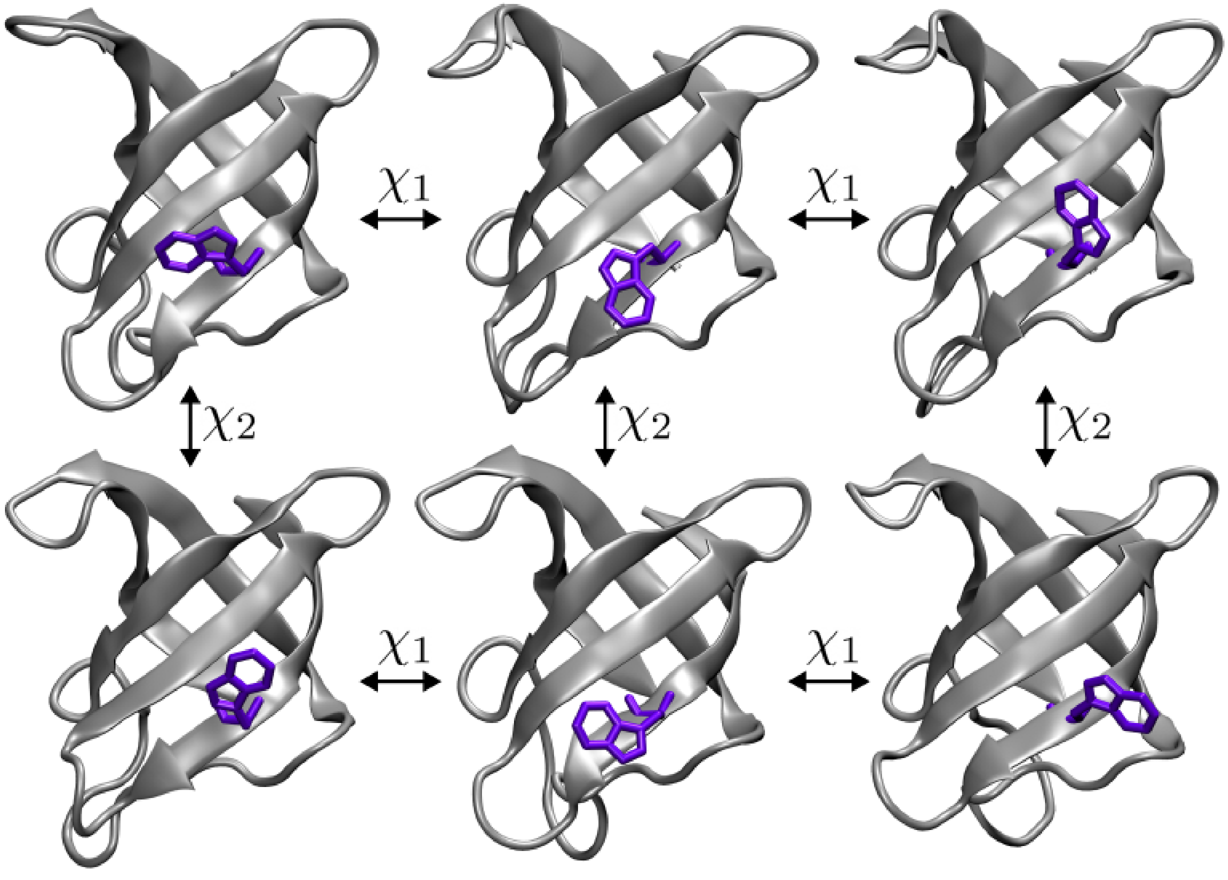


FIG. S1. Representative structures for the six metastable states in Trp8 orientations for cold shock protein 1HZB, with Trp8 highlighted in purple.

To demonstrate the power of our method, we assume no knowledge of Trp8, or any other residues with metastable rotamers, and at every stage of the algorithm, we use all available 125 protein $\chi$ torsional angles. To benchmark thermodynamic weights, we run metadynamics[2] on the same system, but biasing along the Trp8 $\chi_1$ and $\chi_2$ angles so as to generate a PMF in the space. We show that without using any *a priori* information regarding residues that are known to exhibit conformational metastability, we are able to obtain different competing conformations together with their accurate Boltzmann weights. Without using any specific preference towards the $\chi_1$ and $\chi_2$ dihedrals of Trp8, our protocol correctly ranks the 6 metastable states along these two dihedrals.

Further, we also show that our protocol is more general than the focused metadynamics run, and samples conformations along not just Trp8 but along all 53 residues that contain sidechains. To quantify our improvement in generalized sampling, we define sidechain dihedral space as the space of all configurations in all dihedrals, and measure volume explored in this space by an n-dimensional histogram, where n is number of sidechain angles for each residue. We define a measure of ergodicity enhancement as the increase in volume of dihedral configuration space explored. In Figure S3, we see that our protocol not only samples the Trp8 residue conformations comparably to metadynamics directly on that residue, but also samples more extensively for all

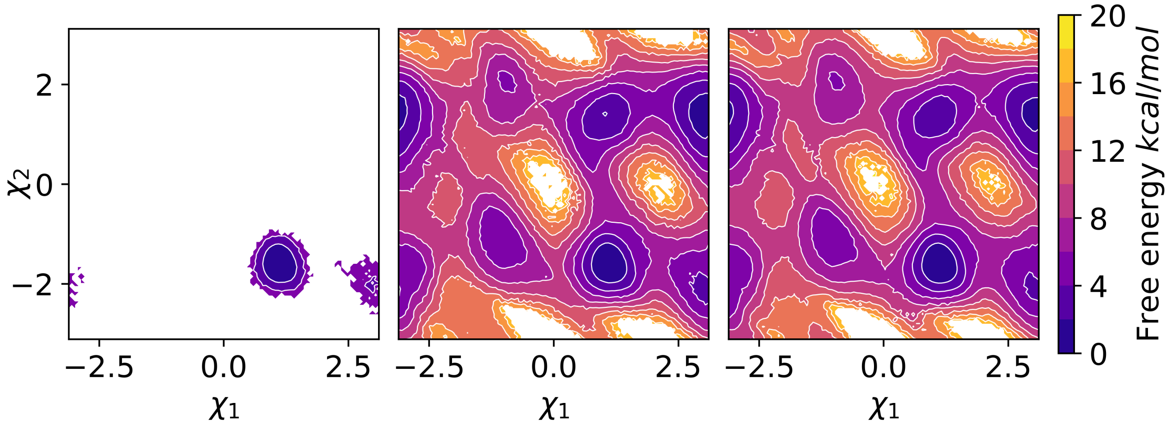


FIG. S2. PMFs along sidechain angles of the Trp8 residue which exhibits six metastable states. In (a) we show calculations from 20 ns long unbiased trajectory initialized from XRD structure. In (b) we show results of metadynamics performed on sidechain angles of the Trp8 residue. In (c) we show results from metadynamics performed using information bottleneck obtained without *a priori* knowledge of Trp8 metastability. Colorbars for all 3 figures are shown with energies in units of $kcal/mol$.

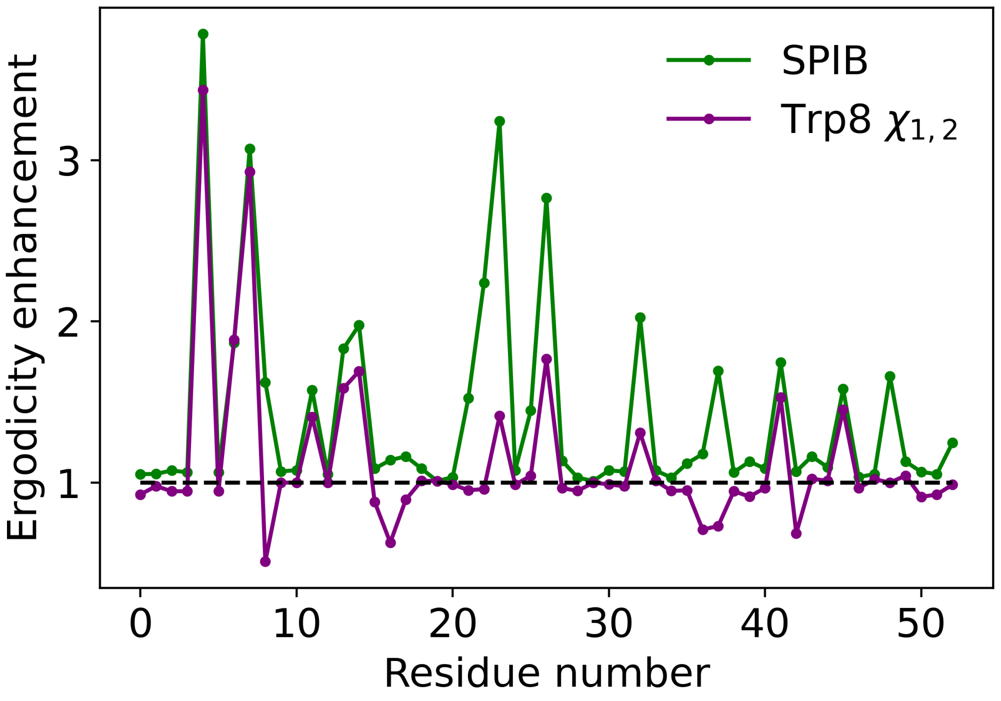


FIG. S3. Ergodicity enhancement in CSP residues. To examine the improvement in sampling of sidechain configurations, we compute the volume sampled in $\chi$ spaces for each residue and plot this for metadynamics performed by (i) directly biasing the sidechain dihedrals $\chi_1$ and $\chi_2$ of the Trp8 sidechains (purple) and (ii) biasing along the information bottleneck our methodology obtains (green). Both volumes are normalized by volume sampled by unbiased trajectories of the same length.

residues with sidechains, which directed metadynamics fails to do.

##### Appendix B: Conformational diversity in SSTR2

As seen in [3], the conformational diversity in GPCRs that is most important for drug design purposes is in interhelical angles and and distances, specifically those of TM6 and TM7. Since we assume no prior information, our collective variables are in terms of $\alpha$-carbon distances of specific residues. However, we can use specific residue distances to establish that interhelical motions are key in our sampling.

Distances with highest weights in our SPIB are PHE92-ASN196 and PHE92-TRP197 (representing TM6-TM7 interactions), GLN126-THR212 (representing TM5-TM6 interactions, and PHE127-ILE209 (representing TM7-TM4) interactions. We also observe significant dihedral motions in helical residues in the ligand binding core of the protein.

**Appendix C: Functionality and structural changes in UBA2**

UBA2, present in the human Rad23 protein [4], and is a 3-helix bundle consisting of 41 residues, where the hydrophobic patch formed between the first and third helix is functionally crucial for binding to ubiquitin, and the constituent helices are extremely stable [5]. Studies show that L355A is primarily a disordered structure[4], [6] with loss of functionality. Our enhanced sampling order parameter is a learnt SPIB, which has non-uniform weights on per-residue Q values, representing a principled approach to combining residue contacts. We use total Q values as a way to characterize the disorder-order separation in UBA2 across both WT and mutant, and to support the stabilization of a partially disordered non-native form to a missense (L355A) mutation. This is a convenient coordinate since it can be decomposed into per-residue Q values and reassembled as a linear combination representation of the SPIB.

Here, we present some more analysis on more structure based collective variables to interpret the mutant's observed loss of functionality. Fig. S4 shows the PMF of both WT and L355A in a 2D space defined by both RMSD and Q-value, showing that our pipeline resolves the structural separation well. The PMF is constructed by reweighting 4 separate trajectories apiece, and we only present the PMF in regions of state space that are sampled by all 4 trajectories so as to minimize error.

Fig. S5 shows fractional helical content for the L355A and WT, i.e. the fraction of residues with backbone torsions corresponding to helices[7] on the Ramachandran plot. These results are presented as a weighted histogram, using weights from our RAVE-based simulations. Fig. S5a shows that about 70 percent of the Wt remains helical with little deviation, while the distribution for L355A is wider and slightly shifted. This corresponds with results in Ref. [8] which show that the protein retains helicity even when partially disordered.

Figs S5 (b,c,d) show fractional helicity per individual helix (H1, H2, H3) as defined in the protein's WT native form. L355A residues from H1 and H3 have a broader distribution compared to the bimodal WT distributions, while the mutation causes a nearly complete loss in H2 helical content. This could suggest that the loss of helical content in H3 may be the cause for the loss of functionality due to mutation, as residues in the H3 along with H1 and loop connecting H1 and

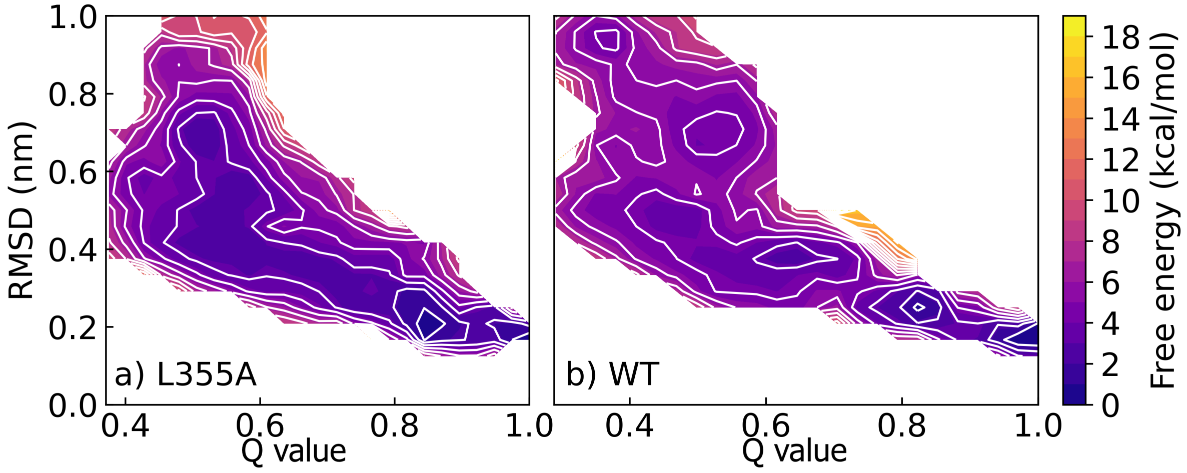


FIG. S4. PMFs plotted on Q value-RMSD space to represent global structural changes found by AF2-RAVE for a) L355A and b) WT. Here, for reference structure, we use an equilibrated structure from classical AlphaFold2.

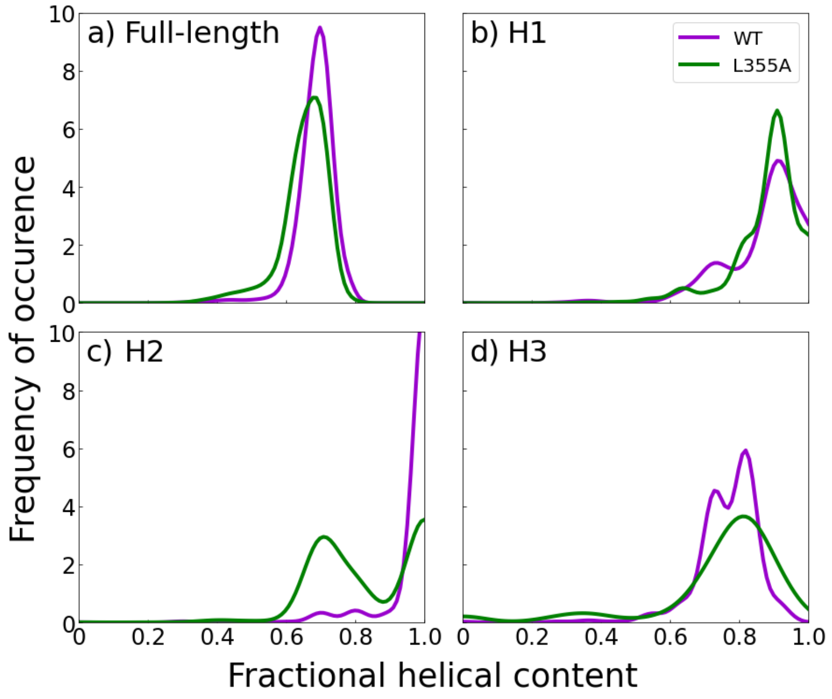


FIG. S5. Fractional helicity for L355A and WT reweighted by metadynamics time-independent weights[2] : a) residues from the full protein structure, b), c), d) residues belonging to helix1 (H1), helix2 (H2) and helix3 (H3), respectively. Helices are defined by structure in the protein's wild type native fold.

H2 form the hydrophobic patch that is crucial for ubiquitin binding.
